# The TPX-Like protein TPXL3, but not TPX2, is the primary activator of α Aurora kinases and is essential for embryogenesis in Arabidopsis

**DOI:** 10.1101/431791

**Authors:** Joanna Boruc, Xingguang Deng, Evelien Mylle, Nienke Besbrugge, Matthias Van Durme, Dmitri Demidov, Eva Dvořák Tomaštíková, Tong-Reen Connie Tan, Michaël Vandorpe, Dominique Eeckhout, Tom Beeckman, Moritz Nowack, Geert De Jaeger, Honghui Lin, Bo Liu, Daniël Van Damme

**Affiliations:** Ghent University, Department of Plant Biotechnology and Bioinformatics, Technologiepark 927, 9052 Ghent, Belgium; VIB Center for Plant Systems Biology, Technologiepark 927, 9052 Ghent, Belgium; Department of Plant Biology, College of Biological Sciences, University of California, Davis, CA 95616, USA; Ministry of Education Key Laboratory for Bio-Resource and Eco-Environment, College of Life Sciences, Sichuan University, Chengdu, Sichuan 610064, China; Leibniz Institute of Plant Genetics and Crop Plant Research (IPK), Gatersleben, Corrensstrasse 3, 06466 Stadt Seeland, Germany; Centre of Plant Structural and Functional Genomics, Institute of Experimental Botany Academy of Sciences of the Czech Republic Slechtitelu 31, CZ-783 71 Olomouc-Holice, Czech Republic; Department of Plant Biology and Linnean Center for Plant Biology, Swedish University of Agricultural Sciences, SE-75007 Uppsala, Sweden.

**Keywords:** Aurora kinase, TPX2, TPX2-Like, spindle formation, Arabidopsis, interactomics, microtubule nucleation

## Abstract

Aurora kinases are key regulators of mitosis. Multicellular eukaryotes generally possess two functionally diverged types. In plants like Arabidopsis, these are termed α versus β Auroras. As the functional specification of Aurora kinases is determined by their specific interaction partners, we initiated interactomics analyses using both α Aurora kinases (AUR1 and AUR2). Proteomics results revealed the TPX2-Like proteins 2 and 3 (TPXL2/3) prominently associating with α Auroras, as did the conserved TPX2 to a lower degree. Like TPX2, TPXL2 and TPXL3 strongly activated AUR1 kinase but exhibited cell cycle-dependent localization differences on microtubule arrays. The separate functions of TPX2 and TPXL2/3 were also suggested by their different influences on AUR1 localization upon ectopic expressions. Furthermore, genetic analyses disclosed that TPXL3, but not TPX2 and TPXL2, acts non-redundantly to secure proper embryo development. In contrast to vertebrates, plants expanded the TPX2 family for both redundant and unique functions among its members.

## Introduction

The evolutionarily conserved Aurora kinases are crucial players in various steps of the eukaryotic cell cycle. Vertebrate Auroras are functionally divided into AUR A and AUR B/C (Carmena and Earnshaw, 2003) based on their differential localization and regulation during the cell cycle. AUR A primarily acts on spindle MTs where it phosphorylates MT-associated proteins (MAPs) in order to enable spindle formation and proper dynamics (Barr and Gergely, 2007). AUR A is targeted to the centrosome and spindle MTs, where it is activated by the MAP TPX2 (for Targeting Protein of the kinesin motor XKLP2) in the frog Xenopus and other vertebrates (Kufer et al., 2002; Wittmann et al., 2000). AUR B, on the other hand, predominantly interacts with the chromosomal passenger complex (CPC, containing the inner centromere protein INCEN-P, Survivin and Borealin). This complex associates with the centromeres in early stages of mitosis and translocates to MTs in the central spindle from anaphase through cytokinesis (reviewed in (Carmena and Earnshaw, 2003)). The spatial compartmentalization and function of the Aurora kinases depends on their interaction partners (Carmena et al., 2009; Li et al., 2015). Remarkably, TPX2-dependent discrimination between AUR A and AUR B relies on a single amino acid close to the catalytic domain and a single amino acid change could functionally convert Aurora A into Aurora B-like kinase (Bayliss et al., 2004; Fu et al., 2009).

TPX2 often mediates interaction of proteins with spindle MTs, and it is widely accepted as an indispensable protein in mitosis (Alfaro-Aco et al., 2017; Gruss et al., 2002). Next to targeting, activating and protecting AUR A from dephosphorylation and degradation, TPX2 makes a critical contribution to MT nucleation inside the mitotic spindle and to chromosome-induced MT assembly (Alfaro-Aco et al., 2017). A more recently reported function of TPX2 is its participation in the DNA damage response (Neumayer et al., 2014). In interphase, TPX2 interacts with importin α and importin β, which shuttle the AUR A-TPX2 complex to the nucleus. High RanGTP levels inside the nucleus mediate dissociation of TPX2 from importins by binding of RanGTP to importin β, thereby driving the accumulation of TPX2 inside the nucleus (Neumayer et al., 2014). In animal cells, a centrosomal pool of TPX2, which aids centrosome separation prior to nuclear envelope breakdown (NEBD), is generated by phosphorylation of TPX2’s nuclear localization signal (NLS) by the NEK9 kinase, that prevents its association with importins (Eibes et al., 2018). Upon NEBD, a high RanGTP concentration, and consequently, high levels of free TPX2, are maintained around the chromosomes due to the association of the RanGEF RCC1 with chromatin. These RanGTP and TPX2 gradients create a positional cue which determines the site of TPX2-mediated MT nucleation (reviewed in (Neumayer et al., 2014)).

In contrast to fungal and animal systems, very little has been learned regarding Aurora-dependent regulation of the cell division cycle as well as on its interaction partners and substrates in flowering plants (reviewed in (Weimer et al., 2016)). Plant Aurora kinases can be classified into α Aurora (AUR1 and AUR2) and β Aurora (AUR3), respectively, in *Arabidopsis thaliana*, based on phylogenetic analysis, differences in subcellular localization and on their differential capacity to complement a weak double mutant in both *Arabidopsis* α Auroras (Demidov et al., 2005; Kawabe et al., 2005). The Arabidopsis α Auroras AUR1 and AUR2 are functionally redundant and associate with the forming spindle and cell plate, whereas β type Aurora localizes to the centromeric region of mitotic chromosomes (Demidov et al., 2005; Kawabe et al., 2005; Van Damme et al., 2011). An *aur1/aur2* double mutant shows defects in division plane orientation mainly during formative cell division in embryogenesis and early stages of lateral root development, suggesting α Aurora’s critical function in establishing cellular asymmetry (Van Damme et al., 2011). Aurora kinases have also been implicated in mitotic and meiotic chromosome segregation in plants (Demidov et al., 2014; Kurihara et al., 2006; Niu et al., 2015) and in securing efficient cell cycle progression through phosphorylation of the MT-bundling protein MAP65-1 (Boruc et al., 2017).

Although a putative INCEN-P homolog termed WYRD, with deviating length and extremely poor sequence conservation to its animal counterpart, has been found (Kirioukhova et al., 2011), it is still unclear whether plants produce a CPC. However, the Arabidopsis genome does contain a clear TPX2 homolog. The canonical TPX2 polypeptide includes an N-terminal hydrophobic Aurora-binding site, a central importin-binding domain, and a C-terminal TPX2 signature MT/kinesin-interacting region, all of which are conserved in the Arabidopsis TPX2 homolog (Vos et al., 2008; Zhang et al., 2017). Arabidopsis AUR1 colocalizes with TPX2 on the spindle MTs, was copurified with TPX2 from Arabidopsis cell cultures and can phosphorylate TPX2 *in vitro* (Petrovska et al., 2012; Petrovska et al., 2013; Tomaštíková et al., 2015). Arabidopsis TPX2 can also bind to *Xenopus* importin α in a RanGTP dependent way (Vos et al., 2008). When antibodies raised against the human TPX2 were injected into the dividing stamen hair cells of the spiderwort *Tradescandia virginiana*, mitosis was blocked because of the inhibition of the formation of the prospindle, which is the bipolar spindle-like MT array formed on the nuclear envelope (NE) at late prophase in plant cells (Vos et al., 2008). Furthermore, *Arabidopsis* TPX2 has MT nucleation capacity *in vitro* and over expression of TPX2 causes ectopic intra-nuclear MT nucleation *in vivo* that is independent of Aurora (Petrovska et al., 2013; Vos et al., 2008) suggesting that TPX2 is essential for bipolar spindle formation in plants, like in animals.

Multiple T-DNA insertional mutations revealed that homozygous *tpx2* mutants did not exhibit obvious cell division or growth phenotypes. This finding implies that the function of canonical TPX2 may be shared with other related proteins. Next to the canonical TPX2, the Arabidopsis genome contains at least eight TPX-Like proteins (TPXLs), of which four bear predicted Aurora-binding motifs (Evrard et al., 2009; Tomaštíková et al., 2015), indicating that the TPX2 family expanded in plants. However, the function of these TPXLs, their connection with plant Aurora kinases, and the potential subfunctionalization of this protein family, remained up to now completely unknown. Here we present functional analyses of TPX2, TPXL2 and TPXL3 in *A. thaliana* that tackle some of these gaps in our knowledge.

## Results

### The TPX-related proteins TPXL2 and TPXL3 are interactors and activators of α Aurora kinases

In the animal kingdom, Aurora kinases play critical roles in multiple cell division processes through interaction with partner proteins and phosphorylation of their substrates (Neumayer et al., 2014). To advance our knowledge on the function of α Auroras in *A. thaliana*, we aimed to identify interactors by tandem affinity purifications (TAPs) coupled with mass spectrometry (MS) analyses, using AUR1 and AUR2 as the bait proteins. We performed two independent TAP tag experiments using both N-terminal (NGS^TEV^), as well as C-terminal (CGS^TEV^) tagged AUR1 and AUR2, followed by mass spectrometry (MS) through tandem matrix-assisted laser desorption/ionization-time of flight (MALDI-TOF/TOF) (Van Leene et al., 2011). Out of these experiments, we identified TPXL2 (TPX-Like2), TPXL3, IMPA1 (α importin 1), and IMPA2 as interactors of both AUR1 and AUR2 (Figure 1A and Source data S1A). Using the improved GS^RHINO^- and GS^YELLOW^-TAP tags fused C-terminally to AUR1 (CGS^RHINO^ and CGS^YELLOW^), combined with the more sensitive LTQ Orbitrap Velos MS (Van Leene et al., 2015), we not only confirmed these four proteins, but also identified seven additional interactors (Source data S1A). TPXL3 was consistently detected in all purifications with either AUR1 or AUR2 as the bait while TPXL2 was recovered in four out of six attempts. In contrast, TPX2 was only found in the Velos MS by using CGS^RHINO^- and CGS^YELLOW^-TAP tagged AUR1. While two α importins, IMPA1 and IMPA2 were detected by both MALDI and Velos with either AUR1 or AUR2, Velos MS revealed IMPA3, IMPA4, IMPA6, and three β importins (karyopherin subunit beta, KPNB1-3). TPXL2 and TPXL3 sequences appear to be very similar to each other and cluster together (Figure 1B, D). We conclude that in contrast to TPX2 proper, which we could only identify using the CGS^rhino^- and CGS^yellow^ tags coupled with the more sensitive LTQ detection, these TPXL proteins were also found in our experiments using the GS^TEV^ tags combined with MALDI detection and therefore can be considered as bona-fide interactors of both α Aurora kinases.

**Figure 1.**
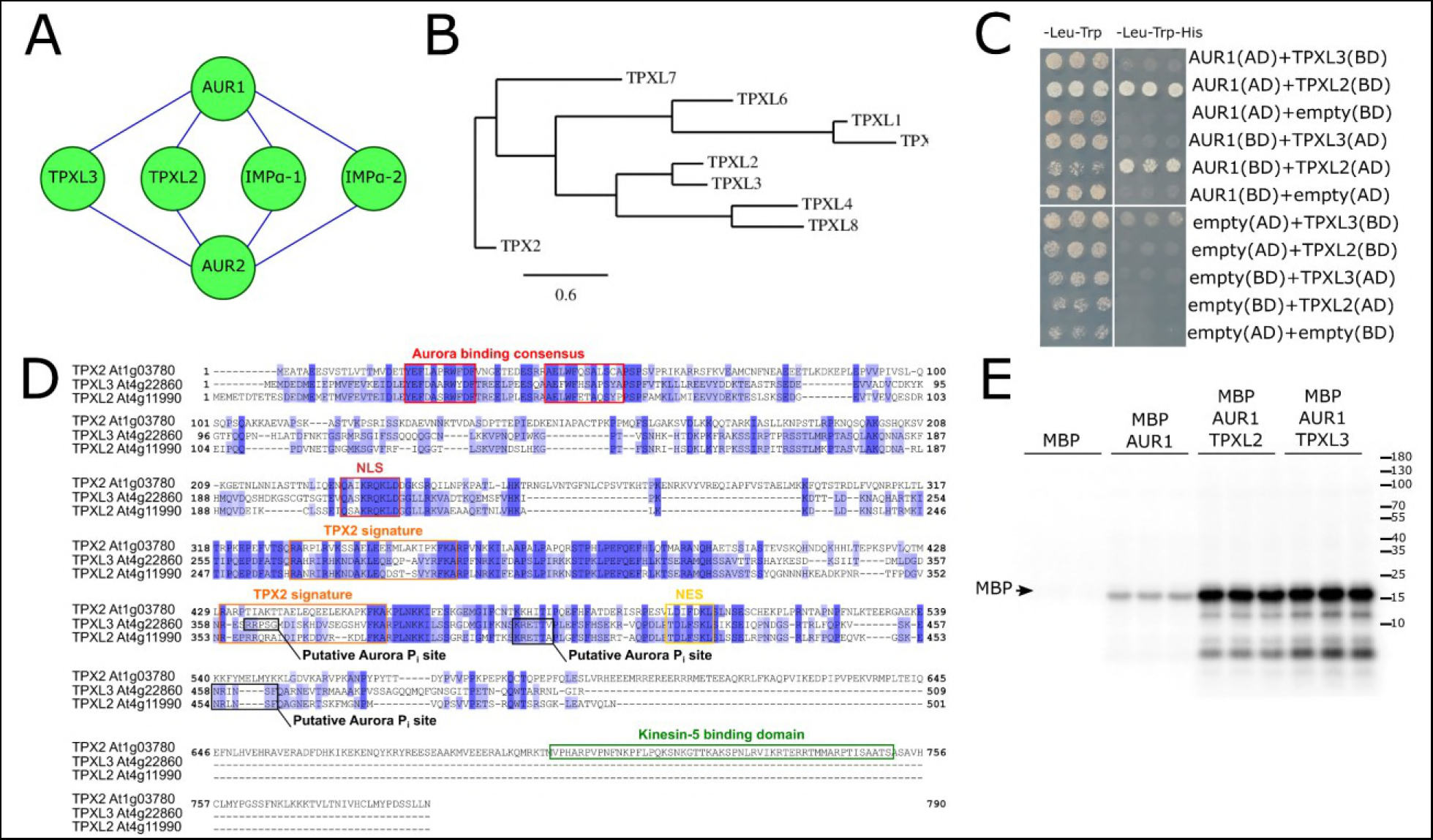
Arabidopsis TPXL2 & TPXL3 are interactors and activators of α Aurora kinases. **A.** Cytoscape representation of the Arabidopsis α Aurora kinases 1 and 2 interactions detected using the Tandem affinity purification (TAP-tag) assay on the Arabidopsis cell suspension culture. Next to two members of the importin protein family, two TPX-Like proteins were co-purified with both Aurora kinases. **B.** Phylogenetic tree of TPX2 & 8 TPX-Like proteins showing the evolutionary relationship among these 9 Arabidopsis proteins. TPXL2 and TPXL3 form a sub-family. **C.** Y2H assay showing confirmation of the interaction between Aurora1 and TPXL2. **D.** Protein sequence alignment of TPX2, TPXL2 & TPXL3 showing that in contrast to TPX2, TPX2L2 and TPXL3 lack the Kinesin-5 binding domain. The Aurora binding consensus motif (red), NLS (light brown), NES (yellow), TPX2 signature (orange), kinesin-5 binding domain (green) and the putative Aurora phosphorylation sites (P_i_; black) are marked with colored boxes. **E.** *In vitro* kinase assay showing that the phosphorylation activity of recombinant Aurora1 is dramatically increased in the presence of the N-terminal TPXL2 or TPXL3 fragment containing the predicted Aurora binding domain (AA 1-100). MBP is myelin basic protein (17kDa).

To take one step further from the TAP results, we tested interactions between AUR1 and both TPXL proteins using yeast two-hybrid analysis. The interaction between TPXL2 and AUR1 could be confirmed in both directions using this system, yet, TPXL3 did not interact with AUR1 in this assay (Figure 1C).

### TPXL2 and TPXL3 share a number of TPX2 motifs

They all bear the N-terminal Aurora-binding domain, to a certain extent the first of the two central TPX2 signature motifs (Vos et al., 2008), a nuclear localization signal (NLS) and a nuclear export signal (NES). Both TPXL2 and TPXL3 however lack the C-terminal Kinesin-5-binding domain typical for TPX2 (Figure 1D). Nevertheless, we did detect 2 or 3 putative Aurora phosphorylation sites within the last 150 amino acid fragment of TPXL2 and TPXL3, respectively (Figure 1D).

The N-terminal part of canonical TPX2, containing the Aurora-binding domain enhances the phosphorylation activity of the kinase (Tomaštíková et al., 2015). To test whether TPXL2 and TPXL3 are also activators of Aurora kinases, we performed *in vitro* kinase assays in the absence and presence of recombinant proteins consisting of the first 100 amino acids of TPXL2 or TPXL3. Whereas recombinant AUR1 is capable of phosphorylating the multipurpose substrate Myelin Binding protein (MBP) *in vitro*, this substrate is hyperphosphorylated in the presence of the N-terminal fragments of TPXL2 or L3, indicating that these TPXL proteins, similar to TPX2, also are potent activators of α-type Aurora kinases (Figure 1E).

### *TPX2, TPXL2* and *TPXL3* expression is correlated with meristematic activity

α Aurora kinase expression is strongly correlated with cell division (Demidov et al., 2005; Van Damme et al., 2011). To test whether the expansion of the plant TPX family could be associated with tissue specific activation of aurora kinases, we analyzed their expression patterns in detail. We therefore generated transcriptional reporter lines for *TPX2, TPXL2*, and *TPXL3* with putative promoter sequences of 2630-bp, 2018-bp, and 1793-bp, respectively. The promoter activities of *TPX2, TPXL2* and *TPXL3* were detected in similar tissues, but overall those of *TPX2* and *TPXL3* seemed stronger than that of *TPXL2* (Supplemental Figure 1). All promoters showed high expression in shoot and root apical meristems, lateral root meristems and in flowers. More specifically, they drove GUS expression in young pistils, ovules and anthers (Supplemental Figure 1). In developing siliques, staining was clearly detected in young seeds for all three promoters although again TPXL2 seemed to have a lower expression compared to the other ones. There was also expression detected in the vasculature of cotyledons and true leaves, again more conspicuous for *TPX2* and *TPXL3.* Expression in the stomatal lineage cells was found only for *promTPXL3* and *promTPX2*, while expression in the flower and silique abscission zone appeared to be more specific for *promTPXL2.*

The observed expression profiles, which correlated clearly with tissues enriched for actively dividing cells, are in agreement with the fact that these proteins are interactors and activators of the mitotic kinases of α Aurora. *PromTPX2, promTPXL2* and *promTPXL3* activity showed clear overlap in dividing root tissues as well as during embryo development, indicating that at least in these tissues, it was unlikely that α Aurora kinases were selectively activated by specific TPX family members.

### TPX2, TPXL2 and TPXL3 interact with AUR1 in planta

To corroborate the interaction between AUR1 and the three TPX2 family proteins *in planta*, AUR1 was transiently co-expressed with either TPX2, TPXL2 or TPXL3 in *Nicotiana benthamiana* leaves. The localization of the individual fluorescent fusions showed that the AUR1-RFP signal was diffuse in the cytoplasm and the nucleus in these epidermal pavement cells (Figure 2A). TPX2-GFP and TPXL3-GFP overall formed nucleus-associated aggregates and filamentous structures, resembling a ball of yarn. In contrast, TPXL2 was more diffuse than filamentous, with the majority of the nuclei showing a pronounced localization close to the NE together with sporadic bright patches (Figure 2A and 2C). Co-expression of AUR1-RFP and GFP fusions of the three TPX family proteins led to distinct relocalization of AUR1. Instead of being cytoplasmic and nuclear, AUR1-RFP became restricted to the nucleus when it was co-expressed with either TPX2-, TPXL2- or TPXL3-GFP (Supplemental Figure 2). Surprisingly, we also observed changes in the nuclear localization of AUR1, TPX2 and TPXL2 when they were expressed simultaneously. When coexpressed with AUR1-RFP, TPX2-GFP mostly lost its filamentous localization, but shared diffused localization with AUR1. Co-expression of AUR1-RFP and TPXL2-GFP on the other hand, altered the localization of both proteins to nucleus-associated filamentous cables (Figure 2B and 2C). TPXL3 retained its filamentous nuclear localization and recruited AUR1 to these cable-like structures. To determine on which side of the NE the observed cable-like structures formed, we used a nuclear envelop marker, RanGAP1-BFP which was co-expressed with AUR1-RFP and either TPXL2-GFP or TPXL3-GFP (Figure 2E). The GFP and RFP signals co-localized with each other and were confined within the RanGAP1-delineated space, showing that these cable-like structures are intra-nuclear.

**Figure 2.**
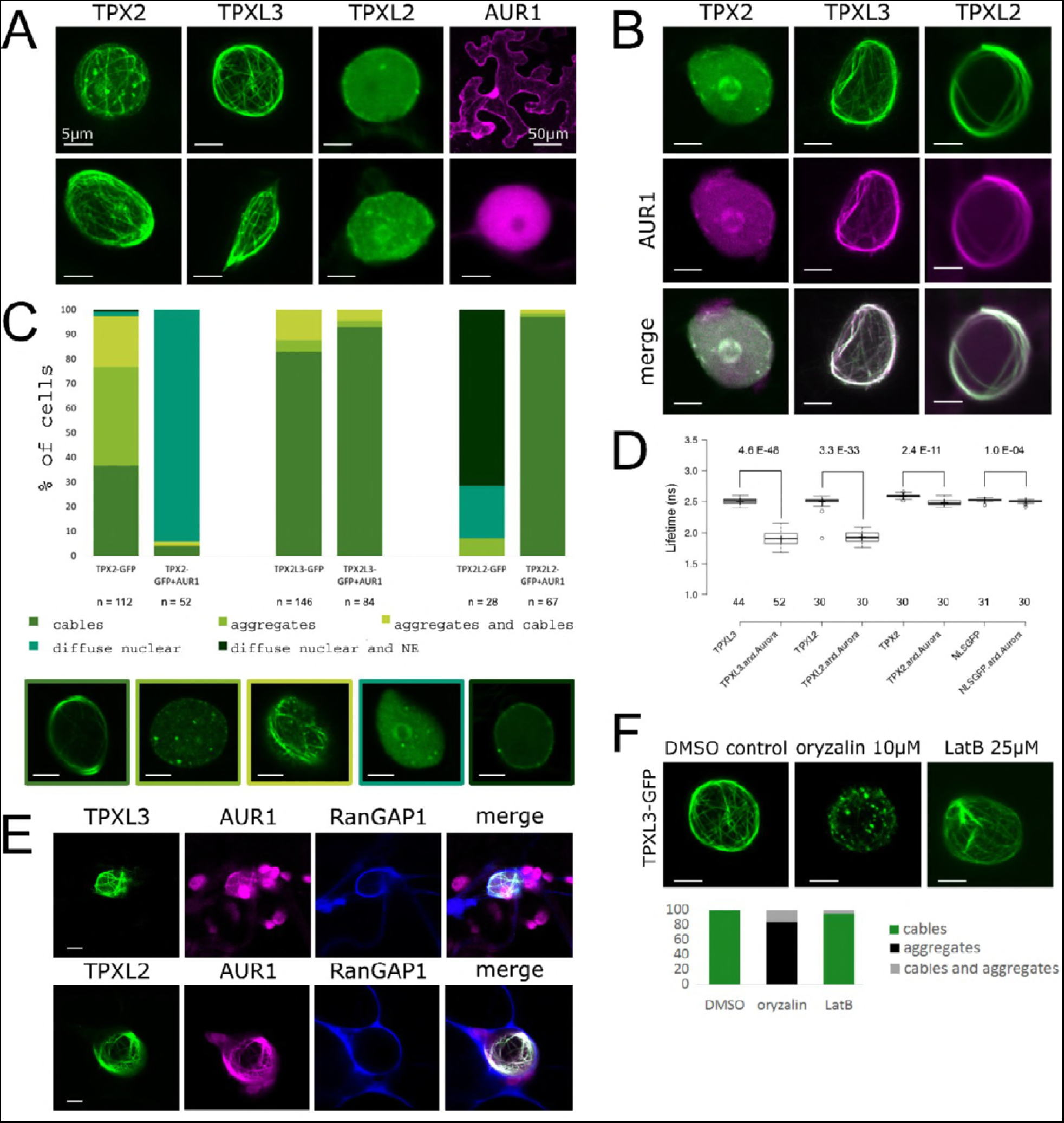
TPX proteins interact with α Aurora 1 in planta and initiate intra-nuclear MT nucleation/polymerization. **A to C.** Representative localizations (A and B) and quantification of the observed localization patterns (C, n equals number of nuclei) for TPX2, TPXL3 and TPXL2 with and without AUR1 in *N. benthamiana* leaf epidermal cells. Single expression of TPX2 and TPXL3, but not TPXL2, causes the formation of intra-nuclear aggregates and cables resembling cytoskeletal filaments. AUR1 is diffuse nuclear and cytoplasmic (A and C). Co-expression of TPX2, TPXL2 and TPXL3 with AUR1 differently affects TPX/L localization. AUR1 has a negative impact on the bundling activity of TPX2, is recruited to the bundles formed by TPXL3 and activates the bundling activity of TPXL2 (B and C). The clear re-localization of AUR1 when combined with high levels of TPXL2 an L3 confirms their interaction. Images below panel C represent the different classes of localizations observed. Left to right: cables (TPX2-GFP); aggregates (TPX2-GFP); aggregates and cables (TPX2-GFP); diffuse nuclear (TPX2-GFP and AUR1); diffuse nuclear and NE (TPX2L2-GFP). **D.** FLIM analysis of co-transformed *N. benthamiana* epidermal cells. Co-expression of TPX-GFP with AUR1-mRFP reduces the donor lifetime and confirms interaction between AUR1 with all three TPX proteins. The reduction in lifetime values is the most pronounced for TPXL2 and TPXL3 (from 2.5ns to 1.9ns); the lifetime decrease of TPX2 is less dramatic (from 2.59 to 2.48). The lifetime of NLS-GFP hardly changes when combined with AUR1 (from 2.53 to 2.51). Numbers represent T-test P values (top) and the number of nuclei analyzed (bottom). **E.** Triple localization of TPXL3 or TPXL2 (green), AUR1(magenta) and the NE marker RanGAP1 (blue) in *N. benthamiana* epidermal cells shows that the cable-like structures form inside the nuclei outlined by RanGAP1. **F.** The intra-nuclear cables marked by the TPX proteins are MTs. TPXL3-GFP localization to the cable-like intra-nuclear structures is sensitive to 1h treatment of 10μM oryzalin (n=25) and changes from intra-nuclear cables into a predominantly aggregated pattern in nuclei when compared to the DMSO control (n=20). Latrunculin B does not affect the localization of TPXL3-GFP (1h 25μM n=21). All scale bars equal 5μm, except for the top right corner image in panel A (50μm).

The intra-nuclear interactions between AUR1 and the TPX proteins were assessed using fluorescence-lifetime imaging microscopy (FLIM) by comparing the lifetime of GFP-tagged TPX2, TPXL2 and TPXL3 in single transformations with those in the presence of AUR1-RFP (Figure 2D). For TPXL2 and TPXL3, the lifetimes decreased from approximately 2.5 ns (single infiltration) to 1.9-ns (double infiltration). This lifetime value drop indicates direct interaction between AUR1 and TPXL2 or TPXL3. The lifetime reduction for TPX2 was much smaller than for the TPX-Likes (from 2.59 to 2.48ns; Figure 2D). As a control, we used NLS-GFP. As expected, the lifetime of NLS-GFP was indistinguishable in the absence or in the presence of AUR1 (from 2.53 to 2.51ns). Although some energy transfer could be observed when both fluorophores were present in the nucleus, the statistical significance differed by several orders of magnitude compared to TPX2 and the TPXL proteins.

Next, we aimed to determine the nature of these cable-like structures using TPXL3-GFP as a representative marker in this *in vivo* experiment. Tobacco leaves infiltrated with the fluorescent fusion were subsequently injected with either oryzalin or Latrunculin B (LatB) in order to depolymerize MTs and actin microfilaments, respectively. Oryzalin reverted the TPXL3-GFP-labeled filaments to distinct nuclear foci. LatB, on the other hand, did not affect the structure of the TPXL3-GFP filaments (Figure 2F). These results indicate that TPXL3, and very likely also TPX2 and TPXL2, associate with intra-nuclear MTs.

### TPX2 is dispensable for α Aurora localization and spindle formation

An earlier report stated that TPX2 was essential for spindle formation and there was no viable homozygous *tpx2* mutant recovered when T-DNA insertional lines were screened (Vos et al., 2008). We aimed to determine the cause of such a lethality by examining community-generated T-DNA lines. To our surprise, homozygous mutants of five different T-DNA insertions in three exons and two introns of the *TPX2* gene were recovered. None of these homozygous lines exhibited any aberrant morphological phenotype as they grew indistinguishably from the Col-0 control plants (Supplemental Figure 3). qRT-PCR analyses revealed that the expression of full-length *TPX2* was absent in *tpx2-1, tpx2-2* and *tpx2-5* mutants (Figure 3B). We also assayed TPX2 expression in the *tpx2-3* and *tpx2-4* mutants with insertions in the 7^th^ and 11^th^ exons, respectively (Figure3A). The RT-PCR result showed both were also likely knock-out mutants as the TPX2 mRNA was undetectable (Figure 3C).

**Figure 3.**
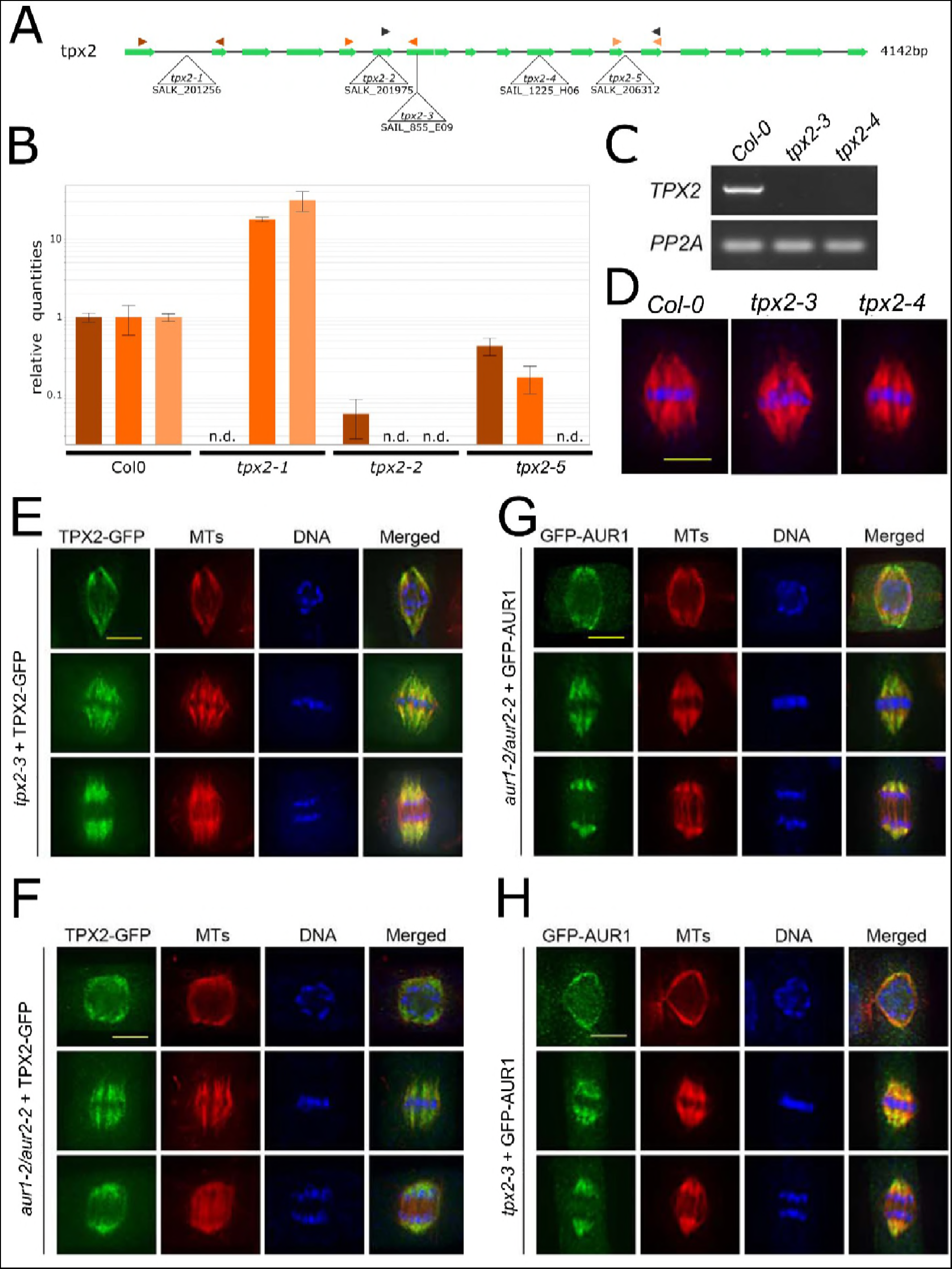
Mutant analysis reveals TPX2 as non-essential. **A.** Schematic overview of the gene model for *TPX2* with indication of the positions of the T-DNA insertion alleles analyzed. Primer pairs used for qPCR and RT-PCR analysis are indicated by color-coded arrowhead pairs. **B.** QPCR analysis of homozygous *tpx2-1, tpx2-2* and *tpx2-5* using the three different primer pairs shown in panel A. The graph shows the absence of full length transcripts over the T-DNA positions (n.d., not detected). **C.** By using the primers marked by the black arrowheads in (A), RT-PCR of homozygous *tpx2-3* and *tpx2-4* showing the absence of transcript compared to WT (Col-0). The constitutively expressed *PP2A* gene was set as the positive control. **D.** Similar spindle MT arrays are formed in metaphase cells in the control (Col-0) and homozygous *tpx2-3* and *tpx2-4* plants. The immunofluorescent images have MTs pseudo-colored in red and DNA in blue. **E and F.** TPX2-GFP localization upon expression in the null *tpx2-3* mutant (E) and in homozygous *aur1-2/aur2-2* double mutant cells (F). Representative cells are at late prophase (top row), metaphase (middle row), and anaphase (bottom row). TPX2-GFP is pseudocolored in green, MTs in red, and DNA in blue. TPX2-GFP decorates the prospindle MTs in prophase and kinetochore fiber MTs at metaphase and anaphase. No obvious difference was detected between the control and *aur1-2/aur2-2* mutant cells. **G and H.** GFP-AUR1 localization in complemented *aur1-2/aur2-2* (G) and in *tpx2-3* mutant cells (H). Representative cells are at late prophase (top row), metaphase (middle row), and anaphase (bottom row). GFP-AUR1 is pseudo-colored in green, MTs in red, and DNA in blue. GFP-AUR1 decorates the prospindle MTs in prophase and kinetochore fiber MTs at metaphase and anaphase. No obvious difference was detected between the control and *tpx2* mutant cells. All scale bars equal 5 μm.

Because TPX2 was implicated in spindle formation (Vos et al., 2008), we examined spindle morphology in the *tpx2-3* and *tpx2-4* mutants when compared to the control cells. Cells of all mutant lines formed mitotic spindles indistinguishable from those in the wild-type control cells. Metaphase spindles of the control, *tpx2-3*, and *tpx2-4* cells had a fusiform appearance with chromosomes perfectly aligned at the metaphase plate (Figure 3D). Therefore, both lines of evidence led to the conclusion that TPX2 is neither essential for spindle assembly, nor the growth and reproduction in *A. thaliana*, a model representing dicotyledous angiosperms.

Considering the fact that the *tpx2* mutants did not show mitotic or developmental, we asked whether TPX2, by analogy to its primary function in animal cells, has a role in targeting plant Aurora kinases to the spindle. We therefore compared the localization of TPX2-GFP and GFP-AUR1, both expressed under the control of their native promotor, in their respective and complementary mutant backgrounds. Among mitotic cells observed by immunolocalization, three representative stages were selected: late prophase (around the time of NEBD when the prospindle could be discerned), metaphase (chromosomes aligned in the middle of a well-established bipolar spindle), and anaphase (shortened kinetochore MT fibers). TPX2-GFP expressed under the control of its native promoter in the *tpx2-3* mutant decorated a fusiform-shaped MT array on the nuclear envelope, (the pro-spindle), in late stages of prophase (top panel, Figure 3E). It associated with spindle MTs, especially kinetochore fiber MTs, and became more pronounced towards spindle poles in both metaphase and anaphase (middle and bottom panel, Figure 3E). As the spindle localization of TPX2 is independent from Aurora A in animal cells (Kufer et al., 2002), we asked whether a similar phenomenon could be observed in plant cells by employing the *aur1-2/aur2-2* double mutant. In the *aur1-2/aur2-2* mutant background, the prospindle was often distorted and did not exhibit an obvious converging morphology (top row, Figure 3F). However, TPX2-GFP still associated with MT bundles around the NE. At both metaphase and anaphase, TPX2-GFP decorated kinetochore fiber MTs, similar to its localization in the control cells (middle and bottom rows, respectively, Figure 3F). Therefore, we concluded that strongly reduced levels of α Aurora do not prevent TPX2 association with spindle MTs *in planta.*

The GFP-AUR1 fusion fully complemented the small dwarf and bushy growth phenotype of the homozygous *aur1-2/aur2-2* double mutant in *A. thaliana* (Supplemental Figure 4), similar to what was reported for the C-terminal fusion (AUR1-GFP) (Van Damme et al., 2011). In the *aur1-2/aur2-2* double mutant background, the GFP-AUR1 fusion first associated with MT bundles in the prospindle (top row, Figure 3G). Then it decorated kinetochore fibers of both metaphase (middle row, Figure 3G) and anaphase spindle (bottom row, Figure 3G). The protein was not associated with MT bundles in the spindle midzone between two sets of sister chromatids (bottom row, Figure 3G). The localization pattern is indistinguishable from that detected with the AUR1-GFP fusion in living cells (Demidov et al., 2005; Van Damme et al., 2011). We then examined GFP-AUR1 targeting in the *tpx2-3* mutant cells. Unlike what has been reported in vertebrates (Kufer et al., 2002), GFP-AUR1 retained its localization pattern in the absence of TPX2 in prophase, metaphase, and anaphase cells (top, middle, and bottom rows, respectively, Figure 3H). Therefore, we infer that α Aurora kinases can associate with spindle MTs independently of the canonical TPX2 protein in *A. thaliana.*

**Figure 4.**
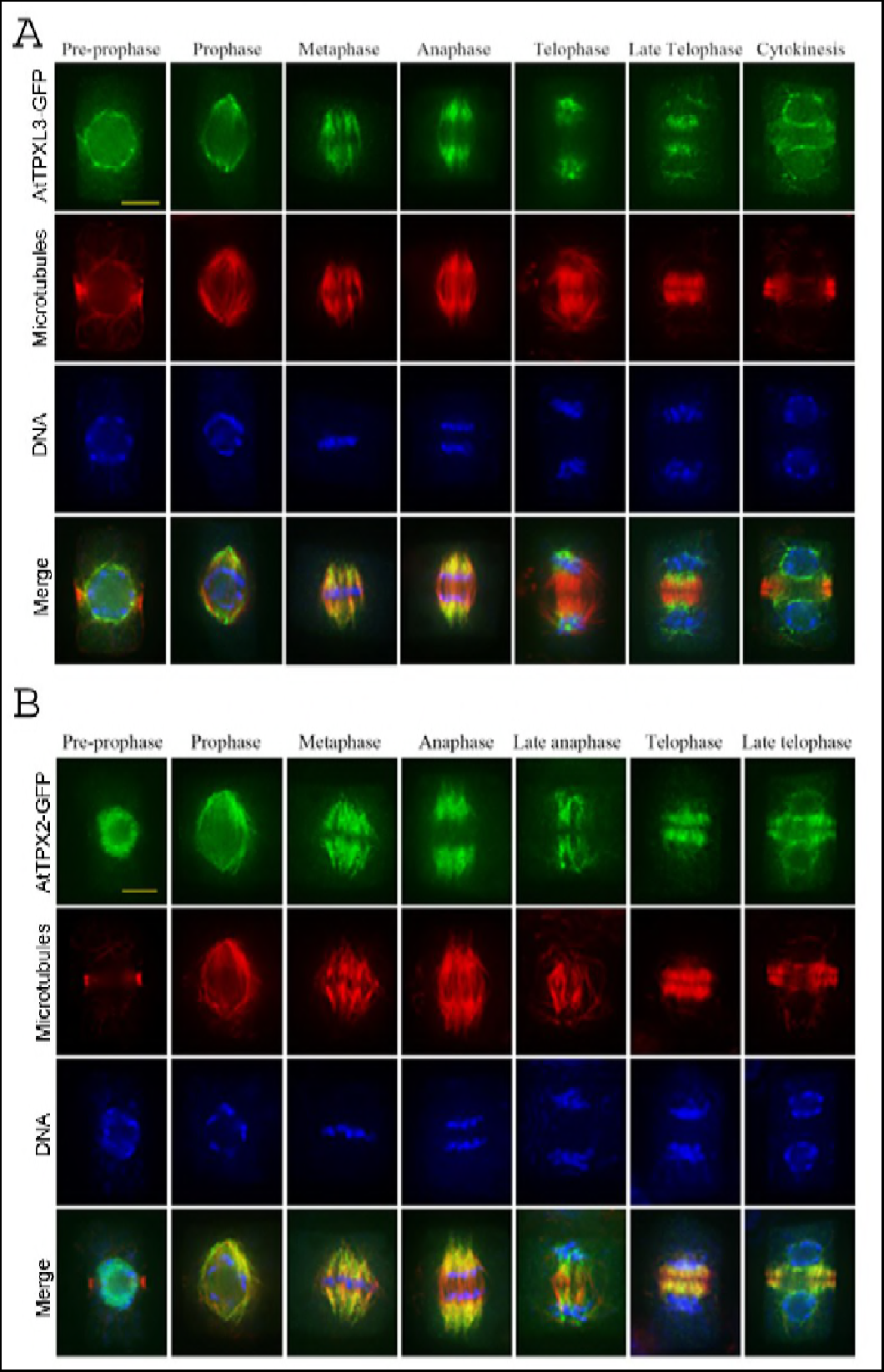
TPX2 and TPXL3 localize differentially in dividing Arabidopsis root cells. **A and B.** TPXL3-GFP (A) and TPX2-GFP (B) localization in dividing Arabidopsis root cells obtained via immunofluorescence with antibodies against GFP (green), microtubules (MTs, red), and DNA (blue) in cells from prophase to telophase. Prior to NEBD, TPXL3 largely accumulates at the NE, while TPX2 is mostly nuclear. Both proteins are not detectable at the preprophase band. Following NEBD, both TPX2 and TPXL3 decorate the two “polar caps” of the prophase spindle as well as spindle MTs and the shortening kinetochore-fibers at anaphase. In late anaphase and telophase, when midzone MTs develop into the two mirrored sets of the early phragmoplast array, TPX2 association with the phragmoplast-forming MTs is much more pronounced than that of TPXL3, which remains heavily enriched at the former spindles poles and subsequently decorates the reformed NE with a strong bias towards the part facing the reforming daughter nuclei. Scale bar, 5 μm.

### The localization pattern of TPXL2 and TPXL3 in dividing cells differs from that of TPX2

Because of the dispensability of TPX2 for mitosis in *A*. *thaliana*, we quested the activities of TPXL2 and TPXL3 in mitotic cells because of their interaction with α Aurora. First, we examined their subcellular localization in tobacco BY-2 cells as a reference system. The two fusion proteins (p35S::TPXL2-GFP and p35S::TPXL3-GFP) showed similar localization patterns in mitosis (Supplemental Figure 5). Prior to cell division, both were detected close to the NE. Immediately prior to NEBD, the TPXL2 and TPXL3 signals appeared to accumulate outside of the NE, co-localizing with MTs marked by the MT-binding domain (MBD) at the prospindle (Supplemental Figure 5). Next, TPXL2 and TPXL3 associated with the metaphase spindle and appeared more prominent on kinetochore MTs (K-fibers) than on the bridging MTs, given the reduced overlap with the RFP-MBD signal at the center of the spindle. In anaphase, both TPXL2 and TPXL3 coincided with the shortening kinetochore fiber MTs. In fact, they became increasingly concentrated toward the spindle poles in both metaphase and anaphase cells. But they clearly avoided the spindle midzone where increased MTs were developing into the bipolar phragmoplast MT array. A striking localization pattern was found during later stages of mitosis when the NE was reforming. Both TPXL2-GFP and TPXL3-GFP fusions heavily localized at the reforming daughter nuclei and eventually formed cage-like patterns. This cell cycle-dependent pattern spatially resembled the cage-like structures observed in the *benthamiana* nuclei. During cytokinesis, both TPXL2-GFP and TPXL3-GFP were clearly absent from the phragmoplast MTs.

Because we understand the potential caveat of ectopic over expression of GFP fusion proteins in a heterologous system, we aimed to determine the localization of TPXL2 and TPXL3 expressed at near endogenous levels *in planta.* To do so, GFP fusion constructs were made using the corresponding genomic fragments including the putative promoters and coding sequences and transformed into wild-type Arabidopsis ecotype Col-0. Because the TPXL2-GFP signal was hardly detectable in our transformed plants, we focused our analysis on TPXL3-GFP. In fact, the localization pattern of TPXL3-GFP in dividing Arabidopsis root cells was comparable to that in BY-2. To gain high spatial resolution, the localization of TPXL3-GFP was determined in fixed cells with MTs and DNA as spatial and temporal references (Figure 4A). Immunostained with an anti-GFP antibody, TPXL3-GFP was detected prominently at the NE in cells with condensed preprophase bands (PPB), but was absent from the PPB, indicating that the protein was selectively associated with certain MTs. Towards late stages of prophase, when NEBD occurs and the PPB dissapears, TPXL3-GFP strongly labeled the polar caps. Similarly to BY-2, kinetochore fiber MTs in both metaphase and anaphase spindles were labeled. After the chromosomes reached the spindle poles at telophase as marked by the nearly complete disappearance of kinetochore fibers, TPXL3 became particularly prominent at the spindle poles where very little MTs were detected (Figure 4A). At this stage, TPXL3 signal was also detected around the chromosome masses, although not as strongly as at the poles. It was, however, largely absent from the central spindle where MTs developed into a bipolar array and formed the phragmoplast. During cytokinesis, TPXL3 accumulated at the reforming NE of the daughter cells.

We also compared the localization of TPXL3-GFP with that of TPX2-GFP in dividing root cells of Arabidopsis (Figure 4B). Although the localization pattern during spindle formation up to anaphase was rather similar for TPXL3 and TPX2, we observed differences at the PPB stage. TPX2 remained primarily nuclear at this point, while TPXL3 accumulated at the NE. Furthermore, during telophase and cytokinesis, and in contrast to TPXL3, TPX2 did mark the phragmoplast MTs but its association with the NE of the daughter cells was not as prominently as that of TPXL3. In conclusion, our localization data in both BY-2 and Arabidopsis cells shows that regulatory mechanisms remain to be characterized that account for the overlaps as well as differences in the targeting of TPX2 as compared to the TPX-Like proteins TPXL2 and TPXL3. These differences mainly concern the capacities to associate with the NE before division and during cytokinesis.

### TPXL3 is essential for plant development

To investigate the functions of *TPXL2* and *TPXL3* in plant development, two independent T-DNA insertional mutations were identified for each gene (Figure 5A). For *TPXL2*, two homozygous mutants were recovered lacking full-length transcript (Figure 5B). Both mutants developed similarly compared to the control plants (Supplemental Figure 3). In contrast to *TPX2* and *TPXL2*, however, we failed to recover homozygous plants for either of the *TPXL3* T-DNA insertional lines. Antibiotic selection revealed that offspring of *tpxl3-1* (SAIL, confirmed by genotyping PCR to be heterozygous for the T-DNA) and offspring of *tpxl3-2* (GABI-Kat, confirmed by genotyping PCR to be heterozygous for the T-DNA), segregated according to a 2:1 ratio of resistant versus sensitive plants (Figure 5C). Reciprocal back-cross experiments to Col-0, using both heterozygous *tpxl3-1* and *tpxl3-2* insertion lines, revealed transmission of the T-DNA via both the male and female, although a decrease in the female T-DNA transmission frequency was observed for *tpxl3-1* (Figure 5C). A close inspection of the developing siliques of the self-pollinated *tpxl3-1* and *tpxl3-2* lines revealed that the mutant siliques were approximately 20% shorter than Col-0 siliques (Col-0, 1.36±0.061(SD) cm, n = 20; *tpxl3-1*, 1.15±0.082(SD) cm, n = 30; *tpxl3-2*, 1.09±0.08(SD) cm, n = 20). These siliques contained a high percentage of aborted ovules and some aberrant seeds (Figure 5D) indicating defects during fertilization and/or during initial embryo development.

**Figure 5.**
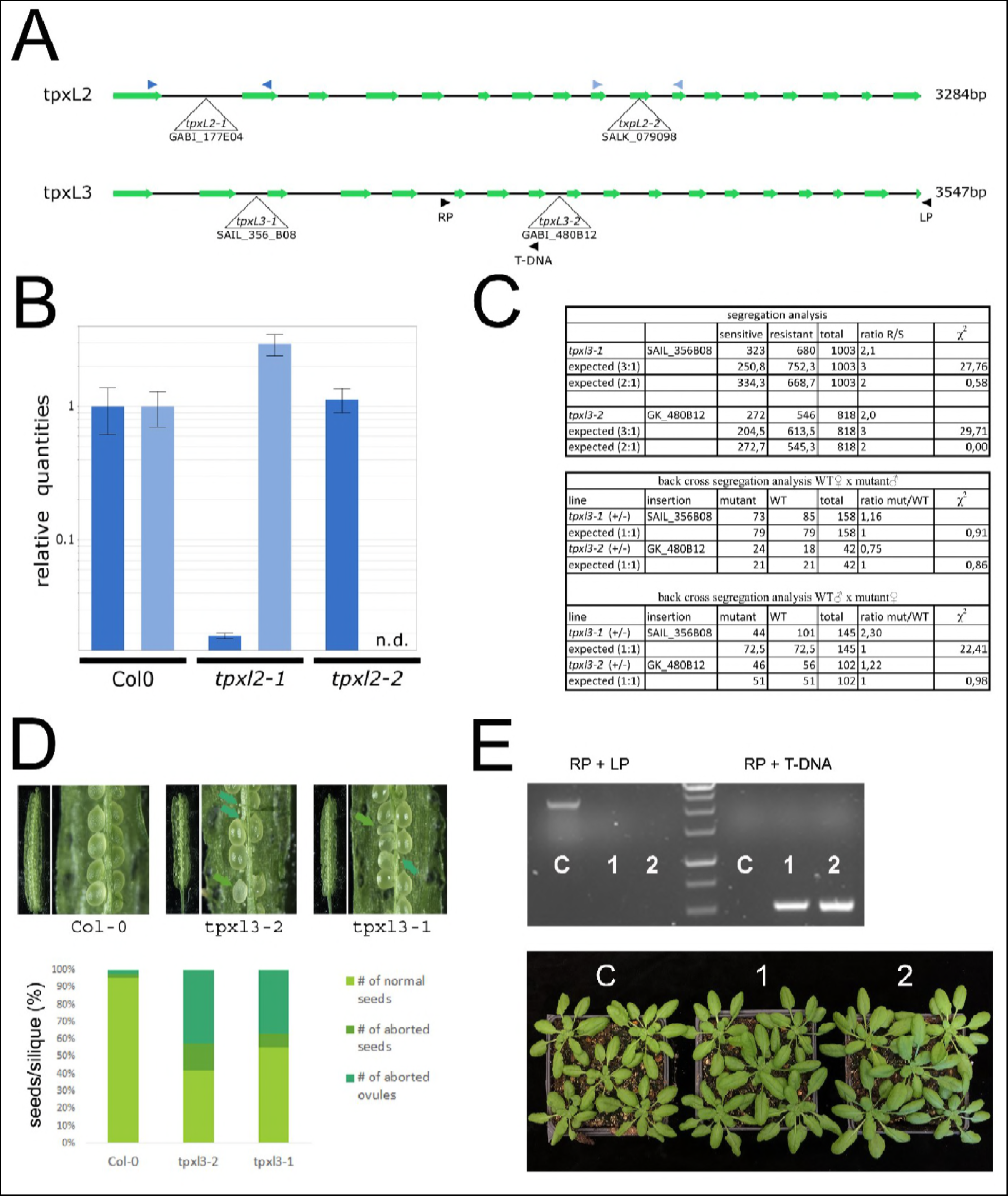
Mutant analysis revealed TPXL2 as non-essential whereas TPXL3 is essential for embryo development. **A.** Schematic overview of the gene models for TPXL2 (top) and TPXL3 (bottom) with indication of the positions of the T-DNA insertion alleles analyzed. Primers used for qPCR and genotyping analysis are indicated by color-coded arrowhead pairs. **B.** QPCR analysis of homozygous *tpxl2* mutants showing absence of full length transcripts over the T-DNA positions (n.d., not detected). **C.** Genotyping analysis for both *TPXL3* insertion lines failed to recover homozygous mutants and back-cross experiments to wild type revealed transmission of the mutant allele via both the male and female gametes, although a reduction in transmission of the *tpxl3-1* mutant allele via the female gametes was observed. **D.** Representative silique pictures and quantification of seed development of selfed *tpxl3-1* (+/−) and *tpxl3-2* (+/−) plants grown together with control plants (Col-0). Both *TPXL3* alleles show a high percentage of aborted ovules and some aborted seeds. **E.** Introducing TPXL3-GFP into *tpxl3-2*(+/−) mutants allowed identification of homozygous *tpxl3-2* mutants (two independent lines are shown; 1 and 2) which develop similarly to the wild type controls (C). Genotyping results revealed absence of WT (LP + RP) fragment (~2.5-kb) and presence of the T-DNA specific fragment (RP + T-DNA; ~0.5-kb) in two independent lines carrying the At4g22860-GFP transgene (1 and 2) in contrast to control plants (C).

To determine whether the lethality was linked to the identified insertion in the *tpxl3-2* mutant and also to test the functionality of the TPXL3-GFP fusion protein described in Figure 4, we examined whether the *tpxl3-2* mutation could be genetically suppressed. Using a primer pair specific for the *TPXL3* locus, we were able to distinguish the native *TPXL3* gene and the *TPXL3-GFP* transgene (Figure 5A, E). Indeed, we recovered two plants with homozygous *tpxl3-2* background when the TPXL3-GFP fusion was expressed. The complemented plants developed similarly to controls (Figure 5E). Taken together, in contrast to *TPX2* and *TPXL2, TPXL3* is an essential gene in Arabidopsis, and the TPXL3-GFP fusion used for determining TPXL3 localization (Figure 4) is functional.

## Discussion

Our results showed that in *A. thaliana*, representing dicotyledonous plants, the expansion of the TPX2 protein family has led to the selection of TPXL3 as an essential MAP for spindle function while the canonical TPX2 has become dispensable. We also found that TPX2 and the two TPXL proteins examined here, are *in vivo* partners of α Aurora kinases, showed overlapping as well as differential localizations during mitosis. Thus our studies opened the door to further investigations of the relationship between TPX2 family proteins and α Aurora kinases as well as how the functions of these MAPs are tied together with mitosis-dependent MT reorganization in plants.

The MAP TPX2, initially identified in *Xenopus*, is required for targeting a plus end-directed motor protein (XKLP2) to the minus ends of the spindle MTs. TPX2 is also necessary for MT-nucleation around the chromosomes in response to a local RanGTP gradient (Gruss et al., 2001; Wittmann et al., 1998; Wittmann et al., 2000). Such TPX2-mediated MT nucleation from the chromosomes is essential for bipolar spindle formation in Hela cells (Gruss et al., 2002). Unlike vertebrates, however, the worm *Caenorhabditis elegans* and the fly *Drosophila melanogaster* lack a clear TPX2 ortholog (Goshima, 2011; Ozlu et al., 2005). Instead, the worm contains a protein harboring the Aurora binding domain of TPX2, but lacking the C-terminal kinesin binding domain. This invertebrate TPXL-1 protein shares with vertebrate TPX2 the capacity to activate Aurora A (Bayliss et al., 2003; Ozlu et al., 2005). More surprisingly, although the fly contains a protein with significant homology to TPX2, it lacks both the Aurora and the kinesin-5 binding domains (Goshima, 2011). The model plant Arabidopsis does contain a clear TPX2 ortholog plus several related TPXL proteins. These TPX-Like proteins differ in the presence or absence of domains characterizing vertebrate TPX2, such as the Aurora binding domain, the TPX2 domain or the Kinesin-5 binding domain (Goshima, 2011; Tomaštíková et al., 2015). In contrast to vertebrates, fly and worm, plants contain both TPX2 and TPXLs and therefore possess an expanded family of TPX2 proteins.

The closely related TPXL2 and TPXL3, especially TPXL3, were identified as the primary interactors of the α Aurora kinases *in vivo* by our MS-based interactomic tests, and the interactions were recapitulated by independent assays. In contrast, TPX2 could only be identified using the more sensitive LTQ Orbitrap Velos. Interaction of AUR1-RFP and TPX2-GFP has previously been reported using co-immunoprecipitation from Arabidopsis culture cells upon co-over expression of both proteins (Petrovska et al., 2012), indicating that the low detection frequency observed in our proteomics results was likely caused by the low abundance of TPX2 in cycling cells of the Arabidopsis culture or possibly weak interaction. These results are also consistent with the finding that TPXL3 but not TPX2 or TPXL2 is essential.

Next to TPXL2 and TPXL3, our TAP analyses using AUR1 as bait revealed several importin α and β proteins, known to facilitate nuclear import of various proteins as was recently shown for the auxin response protein BODENLOS (Herud et al., 2016). It is plausible that importins shuttle TPX2 and the TPXLs, and thereby also the Aurora kinases, to the nucleus. Aurora kinases are nuclear prior to cell division (Van Damme et al., 2011) and interaction between Arabidopsis TPX2 and importins was already shown before (Petrovska et al., 2013; Vos et al., 2008). Moreover, we observed a strong nuclear accumulation of *Arabidopsis* TPX2, TPXL2 and TPXL3 in *N. benthamiana* cells and the nuclear and cytoplasmic localization of AUR1 in *N. benthamiana* shifted to exclusively nuclear upon the over expression of any one of the three TPX(L) proteins in this system.

Similar to TPX2 (Tomaštíková et al., 2015), the N-terminal part of TPXL2 and TPXL3 activates AUR1 kinase activity *in vitro.* The findings suggest that these two proteins not only act as targeting factors of AUR1 but also likely determine the phosphorylation of substrates at specialized localizations. The differences in their localizations, especially between TPX2 and TPXL3 during later stages of the cell cycle and around the time of cell cycle exit, support the notion that different MT-associated proteins may be phosphorylated by AUR1 on different MT arrays. Currently, it is unclear how the localization differences are achieved. Because of the presence of the Kinesin-5-binding site in TPX2 but not TPXLs, TPX2 may specifically require Kinesin-5 for its localization. It would be interesting to test potential interaction(s) between TPX2 and the four isoforms of Kinesin-5 in *A. thaliana.*

Our transient over-expression assays using TPX2, TPXL2, TPXL3 and AUR1 in *N. benthamiana* cells recapitulated the previously reported capacity of TPX2 to localize to or to generate intranuclear MTs in *Arabidopsis* cell cultures (Petrovska et al., 2013). Previously, it was concluded that the formation of those intra-nuclear MT bundles by TPX2 did not require Aurora kinase activity and that they were not linked to a specific cell cycle phase (Petrovska et al., 2013). Here differences in the generation of intra-nuclear MTs are found when different TPX(L) proteins were expressed. We also show that AUR1 affects the capacity of TPX2 and TPXL2 to decorate these MTs. Indeed, whereas TPX2 was capable of forming cage-like structures in the absence of AUR1, this capacity strongly declined in the presence of AUR1. This is in agreement with the observation that in cells with TPX2-decorated MTs, AUR1 signal was very low (Petrovska et al., 2013). For TPXL2, the opposite effect was observed as AUR1 co-expression had a pronounced positive effect on the recruitment of TPXL2 (and AUR1 itself) to these MTs. Taken together, these novel phenomena further support the hypothesis that there are non-overlapping functions among the three proteins.

The oryzalin-induced depolymerization of the TPXL3-positive intra-nuclear cables is in agreement with the published immunolocalization data (Petrovska et al., 2013), and shows that the cables are indeed MTs. It is tempting to speculate that the foci remaining in the presence of oryzalin reflect uncharacterized MT-organizing centers (MTOCs) from which these cables polymerize. Alternatively, these punctae could also represent nuclear pores given the connection between TPXL and importins and the similarity of the pictures with previously published data (Wirthmueller et al., 2015). Although not addressed directly, it seems plausible, given the known function of TPX2 in MT-nucleation (Alfaro-Aco et al., 2017) that the intra-nuclear MT cables are generated as a consequence of ectopic TPX(L) expression and/or increased stabilization due to GFP-tagging. At least for TPX2, the position of the GFP appears to affect protein stability as GFP-TPX2 was reported to disappear already in late anaphase in Arabidopsis and BY-2 cells (Vos et al., 2008), while TPX2-GFP still labels the phragmoplast MTs (Figure 3B). Whether the C-terminal tagging of TPXL2 and TPXL3 also stabilizes these proteins comparable to TPX2 and whether this differs from the N-terminal tagging remains to be established as we failed to successfully express GFP-TPXL2 or GFP-TPXL3 in BY-2 cells (data not shown). Future testing for functionality of the TPX2 fusions will require identification of a mutant phenotype to revert, which will likely require higher order mutant combinations with other TPXL family members.

Nevertheless, the results with TPXL3-GFP in Arabidopsis that we present here show that the C-terminal tagging of TPXL3 does not affect its function as it allowed us to identify plants lacking the native *TPXL3* gene. Following cytokinesis, it is possible that intra-nuclear MTs are formed in Arabidopsis cells similarly to BY-2 cells and that this is a consequence of the extended stability of the TPXL3, which is tolerated by the cells. Alternatively, intra-nuclear cage formation has a specific, yet unknown, function. This notion is supported by the discovery of intra-nuclear microtubule-like structures in *Aesculus hippocastanum L.* by electron microscopy (Barnett et al., 1991). Although speculative at this point, the formation of a MT-cage inside the nucleus might aid to shape the daughter nuclei during NE reformation. On the other hand, it is possible that, similar to what happens during cold treatment, over expression of TPX affects the nuclear pores’ capacity to export tubulin following nuclear envelope reformation (Schwarzerova et al., 2006).

Our mutant analyses using independent T-DNA insertion lines shows that TPX2 and TPXL2 are redundant and dispensable for plant development, whereas TPXL3 is essential. For *TPXL3*, the clear 2:1 segregation ratio and back-cross experiments which showed transfer of the T-DNA via the male and female for both insertion lines, indicate that the mutation does not block development of the gametes, but rather that homozygous mutants are lethal at the embryo stage. The analysis of the siliques of self-pollinated heterozygous *tpxl3-1* and *tpxl3-2* mutants did not reveal clear embryo abortion defects, but showed the presence of apparently unfertilized ovules and a minor fraction of aborted seeds instead which superseded the expected ratio of 25% originating from homozygous lethality. How the presence of apparently unfertilized ovules and seed abortion can be reconciled with the observed segregation ratios pointing to embryo lethality as well as the capability of T-DNA transfer via the female remains enigmatic. One explanation might be that the lack of TPXL3 slows down mitotic progression of the female gametes, leading to asynchrony with the germinating pollen and, therefore, the lack of fertilization. While forced fertilization in the back-cross experiments might overcome this asynchrony, allowing for the observed T-DNA transfer via the female. However, this does not explain the observed segregation ratios for both T-DNA insertion lines, pointing to the equal fitness of the wild type and mutant ovules. Future work is required to pinpoint the cause of the lethality of the *tpxl3* homozygous mutants.

We hypothesize that AUR1/2 proteins, require, next to TPX2, one or more of these TPX-Like proteins as targeting and/or activation factors, simply because the plant Aurora kinases, similar to those in other model systems, will likely rely on other proteins for their defined localizations. The expansion of this protein family likely is associated with the specification of Aurora functions in different aspects of mitosis in a spatiotemporally regulated manner. Our findings also prompt us to hypothesize that TPX2 in plants may be largely devoted to MT nucleation in the spindle apparatus, as a redundant mechanism to those that are governed by the augmin and γ-tubulin complexes.

## Acknowledgements

The authors would like to thank Pavla Binarova for sharing material. Part of this work was supported by grants from the Research Foundation of Flanders (G029013N to D.V.D and T.B) and National Science Foundation in the USA (MCB–1616462 to B.L.). D.D. was supported by the BMBF-funded project „Haplotools“.

## Author Contributions

JB, XD, EM, NB, MVD, DD, ET and MD designed and performed experiments. DE, MN, GDJ, HL, BL and DVD designed experiments, analyzed data and discussed results. JB, BL and DVD wrote the initial draft of the manuscript. JB, XD, MVD, DD, DE, TB, MN, GDJ, BL and DVD contributed to finalizing the paper.

## Supplemental Information

**Supplemental Figure 1.**
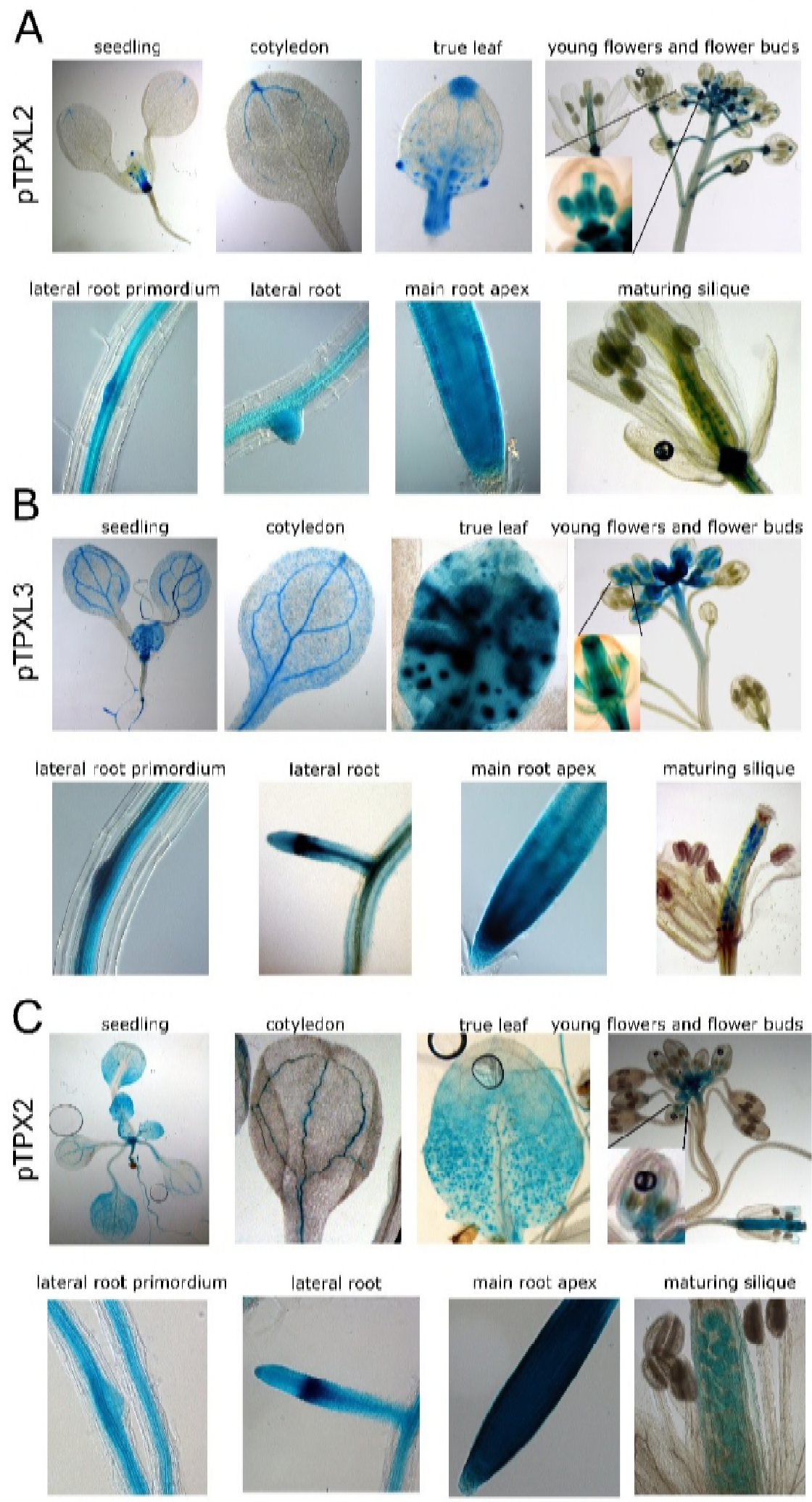
Expression analysis of promTPX2, promTPLXL2 and promTPXL3. Tissue-specific expression of *TPXL2* (A), *TPXL3* (B) and *TPX2* (C) was analyzed using using the respective 5’ genomic sequences driving the GUS transcription. All three putative promoters are active in the shoot and root apical meristems, lateral root primordia, root and cotyledon vascular cells, flowers and developing embryos. Next to these localizations which all represent tissues with high meristematic activity, we also observed promTPXL2 expression in the abscission zone of flowers/siliques and side branches as well as in hydathodes of the cotyledons and the true leaves. PromTPXL3 and promTPX2 were also active in stomata as well as in trichome basal cells in the true leaves. We did not observe clear expression of these promoters in the developing anthers beyond the very early flower stage.

**Supplemental Figure 2.**
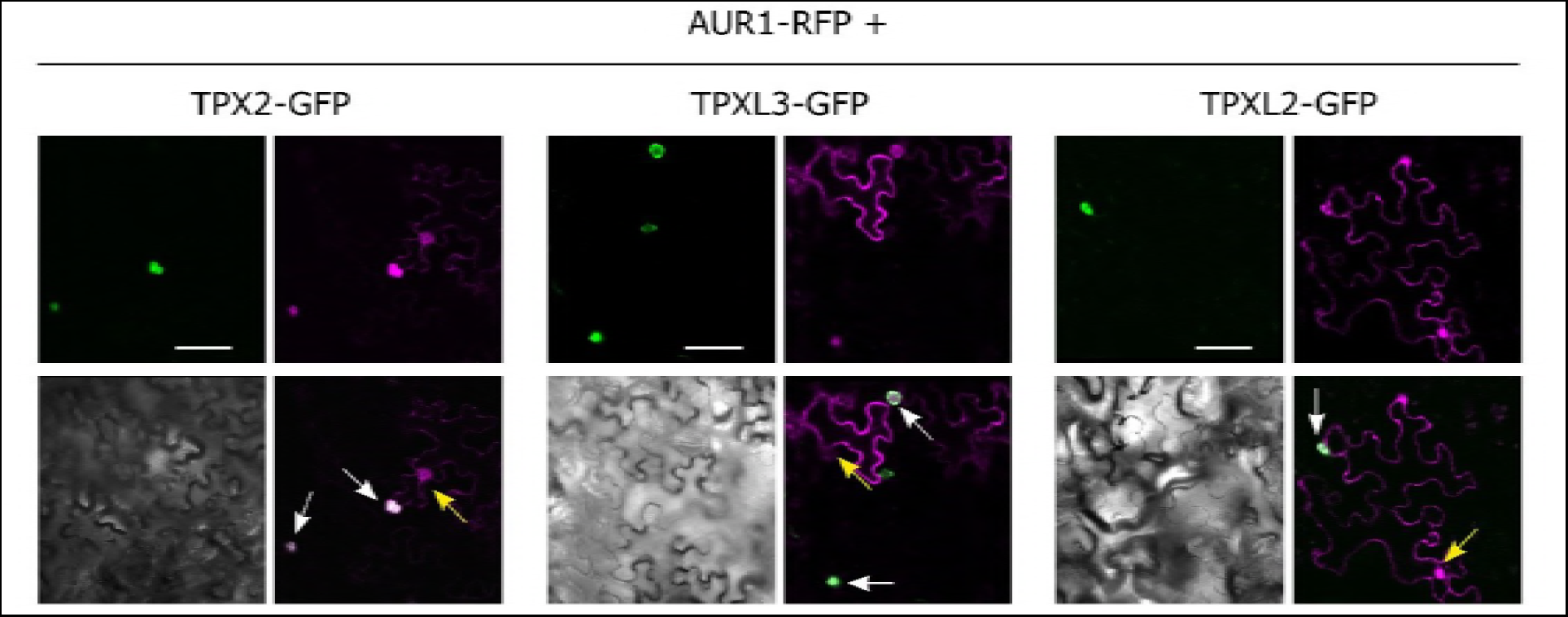
Co-expression of Aurora 1 (magenta) and either TPX2, TPXL2 or TPXL3 (green) restricts the co-localization of Aurora 1 to the nuclear structures in tobacco leaves. White arrows point to the nuclei where both proteins are co-expressed (no cytoplasmic signal for Aur1-RFP), while yellow arrows indicate cells with Aur1-RFP signal alone showing nuclear and cytoplasmic localization of Aurora 1. Scale bars equal 50μm.

**Supplemental Figure 3.**
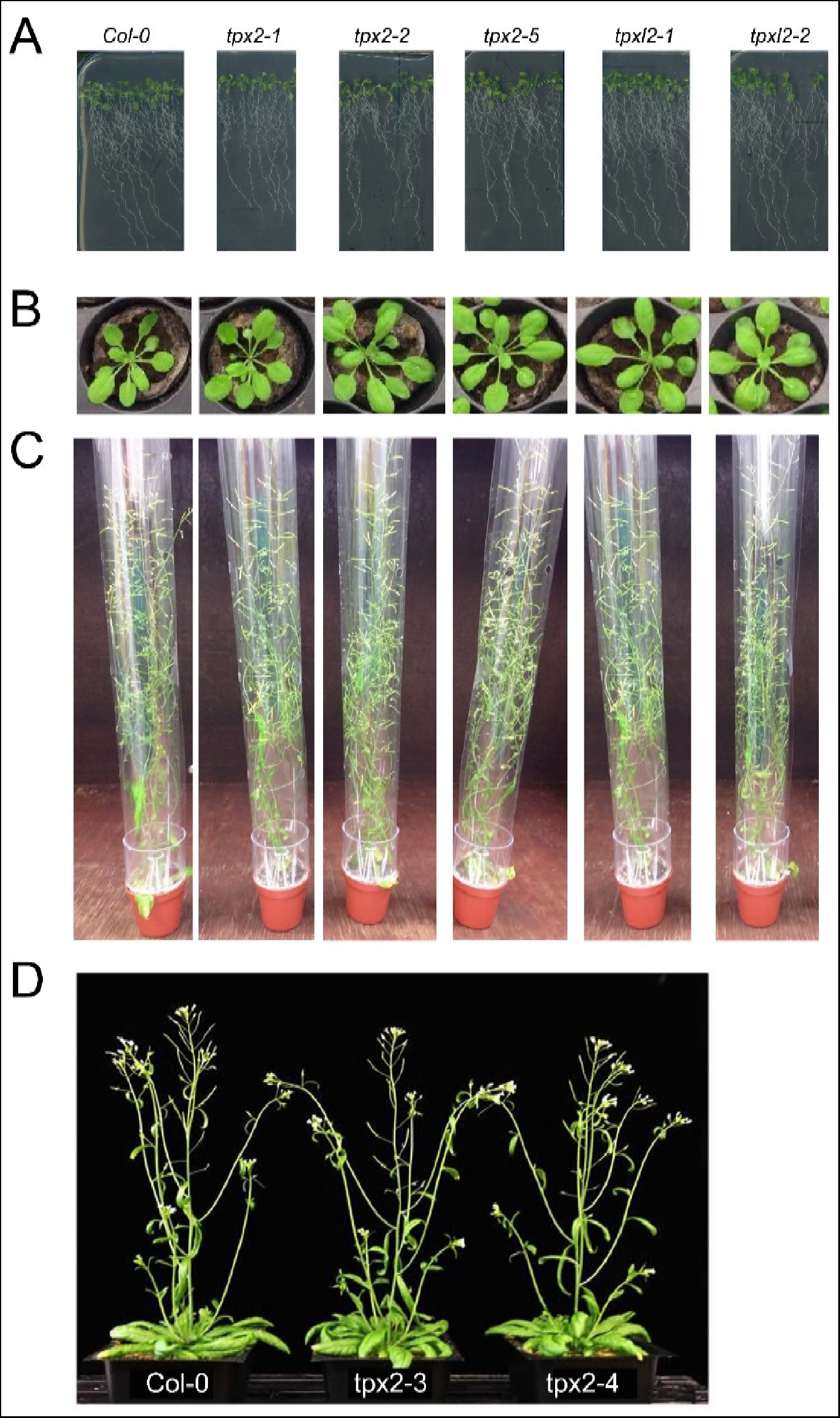
Homozygous *tpx2* and *tpxl2* plants lack any obvious macroscopic mutant phenotype. **A to D.** Representative pictures of young seedlings (A), rosette stage plants (B) and mature plants (C and D) of homozygous mutants in TPX2 and TPXL2 compared to wild type (Col-0). None of the homozygous mutants analyzed for TPX2 and TPXL2 show any obviously developmental defects compared to control plants.

**Supplemental Figure 4.**
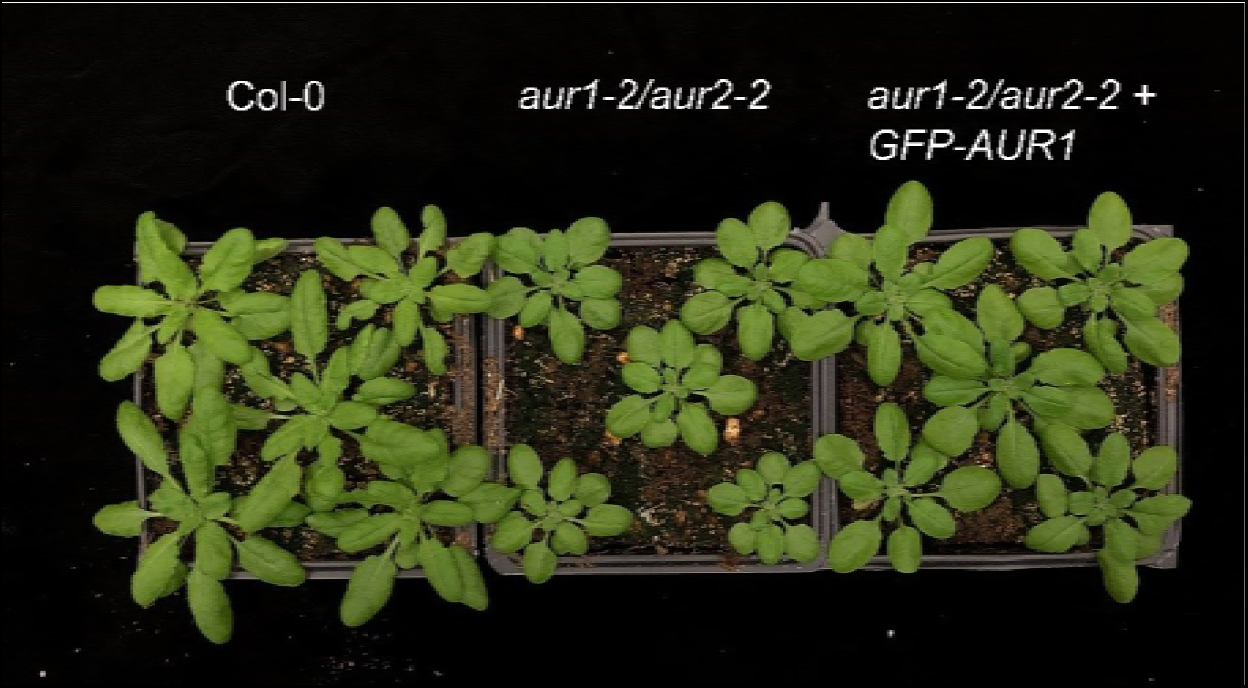
GFP-AUR1 complements the bushy growth phenotype of the *aur1-2/aur2-2* double mutant. Representative 4-week old rosette stage plants of WT (Col-0), *aur1-2/aur2-2* and *aur1-2/aur2-2* with GFP-AUR1 driven by its endogenous promoter. Expression of GFP-AUR1 complements the bushy and condensed growth phenotype of the *aur1-2/aur2-2* double mutant.

**Supplemental Figure 5.**
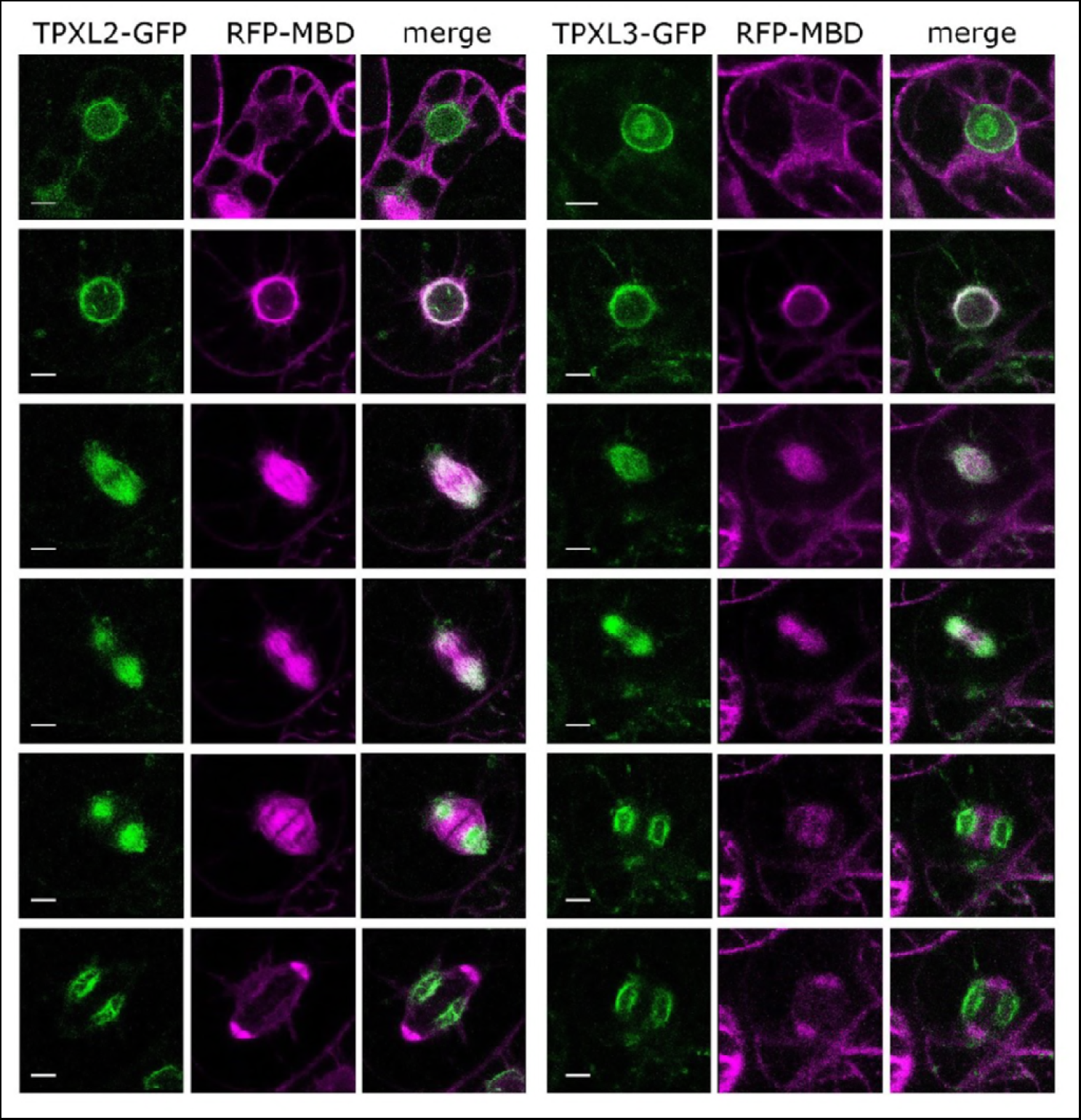
Subcellular localization of TPX2, TPXL2 and TPXL3 in BY-2 cells. Time lapse analyses of dividing BY-2 cells expressing GFP-TPX2 (green, left), TPXL2-GFP (green, middle) and TPXL3-GFP (green, right) combined with RFP-MBD (magenta). P35S::GFP-TPX2 is nuclear before prophase, associates with the forming spindle and follows the separating chromosomes to the spindle poles in anaphase. Spindle MT labeling appears to be restricted to the kinetochore fibers (k-fibers) as no signal is detected on bridging MTs. Following anaphase, the signal disappears. P35S:TPXL2-GFP and P35S:TPXL3-GFP have a similar localization pattern, which differs however from that of GFP-TPX2. Both TPXL2 and L3 predominantly associate with the NE in prophase, label the forming spindle and predominantly k-fibers in anaphase and get recruited to the reforming daughter nuclei forming intra-nuclear cage-like structures in telophase/cytokinesis.

## Supplemental Data files

Source data 1A-1B-1C. List of identified α Aurora interactors using tandem affinity purification.

**Sheet S1A.** Overview of the identified proteins after TAP purification of AUR1 and AUR2 using different TAP tags. Details can be found in sheet S1B for MALDI-TOF/TOF analysis and in sheet S1C for LTQ Orbitrap Velos analysis. Values indicate the number of times a protein was identified in 2 independent TAP purifications. **Sheet S1B**. Protein Identification details obtained with the 4800 MALDI TOF/TOF Proteomics analyzer (AB SCIEX) and the GPS explorer v3.6 (AB SCIEX) software package combined with search engine Mascot version 2.1 (Matrix Science) and database TAIR8. **Sheet S1C**. Protein Identification details obtained with the LTQ Orbitrap Velos (Thermo Fisher Scientific) and Mascot Distiller software (version 2.5.0, Matrix Science) combined with the Mascot search engine (version 2.5.1, Matrix Science) using the Mascot Daemon interface and database TAIRplus.

Supplemental Table1. List of primers used.

## Materials and methods

### Molecular Cloning

#### TAP constructs, Y2H constructs and fluorescent fusions

Full-length open reading frames of TPX2, TPXL2 and TPXL3 were amplified via PCR using primers listed in Supplemental Table 1 and recombined into the pDONR221 entry vector (Invitrogen) by a BP reaction. The LR Gateway reaction resulted in fusions between the open reading frames and GFP, driven by the pr35S in the pK7FWG2 destination vector (Karimi et al., 2007).

Promoters of TPXL2, TPXL3 and TPX2 were cloned into the pDONR221 entry vector and recombined into the pBGWFS7,0 vector in order to generate the promoter::GFP-GUS fusions. The 35S::AUR1-mRFP construct was generated by the Gateway LR recombination using the AUR1 ORF in pDONR207 (Van Damme et al., 2004a) with the pK7RWG2 vector (Karimi et al., 2007). For the yeast two-hybrid assay the genes of interest in pDONR221 were Gateway recombined into pDEST22 (GAL4 AD fusion) and pDEST32 (GAL4 BD fusions) vectors. The 35S::RanGAP1-TagBFP2 construct was generated by Multisite LR reaction into pB7m34GW using pr35S in pDONRP4P1R, RanGAP1 in pDONR221 (Boruc et al., 2015) and TagBFP2 in pDONRP2P3R. To generate the TAP tag constructs for AUR1 and AUR2, TAG-ORF and ORF-TAG constructs, under control of the constitutive cauliflower tobacco mosaic virus 35S promoter, were generated by the Gateway recombination as previously described: N- and C-terminal GS^TEV^ tag (Burckstummer et al., 2006; Van Leene et al., 2008) fusions with AUR1 and AUR2, and C-terminal GS^RHINO^ tag (Van Leene et al., 2015) and GS^YELLOW^ tag (Besbrugge et al., 2018) fusions with AUR1 were constructed.

#### Vectors for the expression of GFP-AUR1 and TPX2-GFP in corresponding mutation backgrounds

The putative promoter region plus the coding sequences were amplified using primers: IV32830F and IV32830R for AUR1, and I03780F and I03780R for TPX2 with Phusion DNA polymerase (Thermo Fisher). The fragments were cloned into pENTR-D/TOPO vector (Thermo Fisher) according to manufacturer’s instruction, resulting the pENTR-AUR1 and pENTR-TPX2, respectively. To produce the GFP-AUR1 fusion construct, the EGFP sequence was amplified using primers of GFP-AUR1F and GFP-AUR1R. The amplified fragment was cloned into linearized pENTR-AUR1 plasmid via the Gibson Assembly method (NEB) to give rise to pENTR-GFP-AUR1. The final expression constructs were made by having pENTR-GFP-AUR1 and pENTR-TPX2 recombined with pGWB10 and pGWB4 (Nakagawa et al., 2007) using LR clonase (Thermo Fisher). These constructs were introduced into the *aur1-2/aur2-2* and *tpx2-1* mutants by Agrobacterium-mediated DNA transformation via the standard floral dipping method.

### Plant Growth and Transformation

Both the wild-type control, and mutants are in the *Arabidopsis thaliana* ecotype Columbia-0 (Col-0) background. All lines (except SAIL_855_E09 and SAIL_1225_H06) were grown under standard growth conditions at the Center for Plant Systems Biology in Ghent in continuous light on vertical plates containing half-strength Murashige and Skoog (MS) medium supplemented with 8 g/L plant tissue culture agar and 1% sucrose. SAIL_855_E09 and SAIL_1225_H06, together with their control were grown at 21°C with 16-hr light and 8-hr dark in growth chambers at the Controlled Environmental Facility on the campus of the University of California in Davis.

Wild-type Col-0 and mutant plants were transformed using the floral dip protocol (Clough and Bent, 1998). The Arabidopsis mutant lines *tpxl3-2* GABI_480B12 and *tpxl2-1* GABI_177E04 were acquired from the Gabi-Kat collection (Kleinboelting et al., 2012). Out of the 40 seeds received from for the *tpxl3-2* mutant, only 8 seeds germinated. Out of these 8 plants, 3 were identified as having the T-DNA (line 3, line 4 and line 7) and we continued working with line 7.

The Arabidopsis mutant lines *tpxl2-2* SALK_079098, *tpxl3-1* SAIL_356_B08, *tpx2-1* SALK_201256, *tpx2-2* SALK_201975 and *tpx2-5* SALK_206312 were acquired from the Arabidopsis Biological Resource Center (ABRC) located at Ohio State University in Columbus, Ohio. Homozygous *tpx2-3* and *tpx2-4* mutant plants were isolated from the seed pools of SAIL_855_E09 (CS838303) and SAIL_1225_H06 (CS844904) lines, respectively, ordered from the ABRC. For *tpxl3-1*, 7 plants were genotyped and 3 were positive for the T-DNA band (line 1, line 4 and line 7). We continued working with those 3 lines for the segregation analysis. The *aur1-2/aur2-2* double homozygous mutant was reported previously (Van Damme et al., 2011).

T-DNA transfer in the reciprocal back-cross experiments of heterozygous *tpxl3-1* and *tpxl3-2* mutants was quantified by antibiotic resistance associated with the SAIL and Gabi KAT T-DNA insertions of the offspring seedlings. Deviations from the expected theoretical segregation ratio for both selfed and crossed offspring seedlings was assessed via Chi square testing of observed versus theoretical values (3:1 and 2:1).

Wild-type tobacco (*Nicotiana benthamiana*) plants were grown under a normal light regime (14 h of light, 10 h of darkness) at 25°C and 70% relative humidity. Tobacco infiltration with Agrobacterium tumefaciens strain C58 was performed as described previously (Boruc et al., 2010). However, in addition, co-expression of the p19 protein from tomato bushy stunt virus was used for suppression of transgene silencing (Voinnet et al., 2000). The quantification of the *benthamiana* localization data shown in Figure 2C is the combined result of 3-8 independent transformation events.

Stable tobacco BY-2 cell culture (*Nicotiana tabacum* ‘Bright Yellow-2’) transformation was carried out as described before (Geelen and Inze, 2001). For the co-localization analysis, BY-2 cells were first transformed with the 35S::RFP-MBD (Van Damme et al., 2004) construct, selected on hygromycin and then transformed again and screened for the presence of the kanamycin-resistant GFP-fusion constructs. Cells derived from several independently transformed calli were imaged for each construct.

### Mutant genotyping

The *tpx2-1, tpx2-2, tpx2-5, tpxl2-1, tpxl2-2, tpxl3-1* and *tpxl3-2* mutations were identified using the genotyping primers listed in Supplemental Table 1 by combining either the LP or RP with the T-DNA specific primer of the SALK, SAIL or GK collection. The *tpx2-3* mutation was detected by PCR using 838303RP and GLB3 and *tpx2-4* by using 844902RP and GLB3. Gene-specific primer pairs spanning the T-DNA insertions for *tpx2-3* were 838303LP and 838303RP and a combination of 844902LP and 844902RP for *tpx2-4.*

### Histochemical Analysis

Seedlings or flowering stems were stained in multi-well plates (Falcon 3043; Becton Dickinson). GUS assays were performed as described previously (Beeckman and Engler, 1994). Briefly, samples were fixed in ice-cold 80% acetone, washed 3 times in the phosphate buffer and stained for 3–5 hours at 37°C. Next, 3 washes in the phosphate buffer were followed by immersion in 100% ethanol. Samples mounted in lactic acid were observed and photographed with a stereomicroscope (Olympus BX51 microscope) or with a differential interference contrast microscope (Leica).

### Tandem affinity purification (TAP) tag assay

Transformation of Arabidopsis cell suspension cultures was performed by Agrobacterium cocultivation and transgenic culture regeneration. TAP purification of protein complexes, further processing, mass spectrometry analysis, data analysis and background filtering was done as described before: GS^TEV^ purifications were analyzed on a 4800 MALDI TOF/TOF Proteomics Analyzer (AB SCIEX) ((Van Leene et al., 2011), Source data S1A and S1B for details), GS^RHINO^ and GS^YELLOW^ purifications were analyzed on a LTQ Orbitrap Velos (Thermo Scientific) as described before ((Besbrugge et al., 2018; Van Leene et al., 2015) Source data S1A and S1C for details).

### Yeast two-hybrid (Y2H) assay

For the yeast two-hybrid assay, plasmids encoding the baits (pDEST32) and preys (pDEST22) were transformed into the yeast strain PJ69-4A (MATa; trp1-901, leu2-3,112, ura3-52, his3-200, gal4D, gal80D, LYS2TGAL1-HIS3, GAL2-ADE2, met2TGAL7-lacZ) and prey by the LiAc method (Gietz et al., 1992). Co-transformed yeast cells were selected on synthetic dextrose (SD) plates without Leu (pDEST32) and without Trp (pDEST22). Interactions between proteins were scored by transferring the colonies on the SD medium lacking Leu, Trp, His and Ade and incubated at 28°C for 2–3 days. The results shown represent three biological repeats and in total the experiment has been performed twice.

### Plasmid Construction for Expression of Recombinant Proteins in *Escherichia coli*

Coding sequences of the N-terminal putative Aurora binding domain (AA 1-100) of AtTPXL2 and AtTPXL3 were cloned as described previously (Petrovska et al., 2012) using primers listed in Supplemental Table 1 (TPXL2for – TPXL2rev and TPXL3for – TPXL3rev). For expression of recombinant 6His-tagged protein, *At*TPXL2-and *At*TPXL3 fragments were subcloned into the Gateway expression vector pET55DEST (Novagen, Madison, WI, USA) and *At*Aurora1 was cloned as described previously (Demidov et al., 2009).

### Production of recombinant proteins

GST-Aurora1 was expressed in *E. coli* C-43 strain and purified under native conditions as described before (Tomaštíková et al., 2015). Aurora binding domains of TPXL2 and TPXL3 proteins were expressed in *E.coli* BL-21 and purified under denaturating conditions according to Demidov et al., 2005 (Demidov et al., 2005).

### *In vitro* kinase assay

Purified recombinant proteins were desalted in kinase buffer using 7K MWCO Zeba Spin Columns (Thermo Scientific) and processed as described in Tomastikova et al., 2015 (Tomaštíková et al., 2015). Briefly, the AtAurora1 alone (as control) or with the same amount of TPX were incubated at 30°C, 30 min with 0.5 × kinase buffer and 0.1 mM ATP for activation of the kinases. Subsequently, [32P]ATP and substrate (myelin basic protein (MBP) ~10 μg) were added and incubated for additional 60 min at 30°C, as described in Tomastikova et al., 2015.

### Immunofluorescence microscopy

Root meristematic cells were processed for immunofluorescence microscopy as described previously (Lee and Liu, 2000). A rabbit anti-GFP antibody and the DM1A mouse monoclonal anti-α-tubulin antibody (both from Sigma Aldrich) were used for the detection of GFP fusion proteins and tubulin/microtubules in fixed root cells.

### Sequence alignment and Phylogenetic Analyses

The following reference sequences of proteins were used to generate the protein sequence alignment and the phylogenetic tree in Figure 1: AT1G03780 (TPX2), AT3G01015 (TPXL1), AT4G11990 (TPXL2), AT4G22860 (TPXL3), AT5G07170 (TPXL4), AT5G15510 (TPXL5), AT5G37478 (TPXL6), AT5G44270 (TPXL7), AT5G62240 (TPXL8). Protein sequence alignment was performed using Clustal Omega (EMBL-EBI) and processed with the Jalview program (Waterhouse et al., 2009). The phylogenetic tree was built using the phylogeny tool (www.phylogeny.fr). The tool uses the MUSCLE multiple sequence alignment and PhyML for the construction of phylogenetic trees. TreeDyn viewer was used to visualize the tree (Dereeper et al., 2010; Dereeper et al., 2008). The tree was rooted on TPX2.

### Q-PCR Expression Analysis

Seven day old seedlings were frozen in liquid nitrogen and ground. RNA was extracted using standard Trizol-chloroform extraction and genomic DNA was removed by DNAse treatment. Reverse transcription was performed using the Promega ImProm-II reverse transcription system according to manufacturer’s instructions. QPCR was performed on the LC480 (Roche) and results were analyzed using Qbase+. Primer sets spanning the T-DNA positions for *tpx2-1, tpx2-2, tpx2-5, tpxl2-1* and *tpxl2-2* as well as normalization primers (CDKA and Actin4) are listed in Supplemental Table 1. The values presented are the average of three technical repeats and the experiment has been performed twice with similar results starting from independent seedlings and independent RNA extraction.

### RNA extraction and reverse transcription PCR (SAIL_855_E09 and SAIL_1225_H06)

Total RNA was isolated from 3-days-old seedlings using PureLink RNA Mini Kit (Invitrogen) according to manufacturer’s instruction. Reverse transcription was performed with iScript cDNA Synthesis Kit (Bio-Rad). The primers used for RT-PCR were TPX2F and TPX2R. The PP2A transcript was used as a positive control as described previously (Czechowski et al., 2005). PCR products resulted from 35 amplification cycles were analyzed by agarose gel electrophoresis.

### Confocal Laser Scanning Microscopy

Fluorescence was analyzed with an inverted confocal microscope FluoView FV1000 (Olympus), equipped with a 60× water-corrected objective (n.a. 1.2); or an LSM710 laser scanning confocal module mounted on an inverted Axiovert system (Carl Zeiss). Fluorescence was imaged in a multichannel setting with 488− and 543-nm excitation light for GFP and RFP excitation, respectively. Emission fluorescence on the Olympus system was captured in the frame-scanning mode via 500− to 550-nm and 560− to 660nm band pass emission windows for GFP and mRFP fluorescence, respectively. Imaging on the LSM710 used 40X C-Plan (water) or 63X Plan-Apo (oil) objectives, and the GFP signal was excited by 488-nm argon laser, and images were acquired using the ZEN software package attached to the confocal system. Confocal images were processed with ImageJ (www.imagej.nih.gov/ij).

### Foster resonance energy transfer-fluorescence lifetime microscopy (FRET-FLIM)

The donor fluorescence lifetime (FLIM) was determined by time-correlated single-photon counting (TCSPC) in tobacco epidermal cells transiently transfected with the proteins of interest fused to either eGFP (donor) or mRFP (acceptor).

Images were collected using an LSM Upgrade Kit (PicoQuant) attached to a Fluoview FV-1000 (Olympus) confocal microscope equipped with a Super Apochromat 60x UPLSAPO water immersion objective (NA 1.2) and a TCSPC module (Timeharp 200; PicoQuant). A pulsed picosecond diode laser (PDL 800-B; PicoQuant) with an output wavelength of 440 nm at a repetition rate of 40 MHz was used for donor fluorescence excitation. Laser power was adjusted to avoid average photon counting rates exceeding 10000 photons/s to prevent pulse pile up. Samples were scanned continuously for 1 min to obtain appropriate photon numbers for reliable statistics for the fluorescence decays. A dichroic mirror DM 458/515 and a band pass filter BD520/32-25 were used to detect the emitted photons using a Single Photon Avalanche Photodiode (SPAD; PicoQuant). Fluorescence lifetimes were calculated using the SymPhoTime software package (v5.3.2.2; PicoQuant). Selected areas of the images corresponding to single nuclei (n ≥ 30 cells from several independent transformations) were fitted by either a single-mono-exponential fitting for a donor alone sample or a bi-exponential fitting for a combined donor-acceptor sample including the measured IRF (instrument response function). The measured IRF was determined using an erythrosine B solution as described for the samples but with a lower count rate (1000 photons/s). The lifetimes τ for a series of measurements were presented in a boxplot showing the median (center lines), mean, standard deviation and the outliers (dots). Significance between donor alone and donor with acceptor samples was checked using Student's T-test, assuming a twotailed distribution and unequal variance.

### Primers used

All primers used in this study can be found in Supplemental Table 1.

### Quantification

Box plot graphs were generated using the BoxplotR web tool (Spitzer et al., 2014).

### Accession Numbers

The Arabidopsis Information Resource (TAIR) locus identifiers for the genes mentioned in this study are At1g03780 for TPX2, At4g11990 for TPXL2, At4g22860 for TPXL3, At4g32830 for AUR1, and At2g25880 for AUR2.

